# ME1 Programs Latent Effector Capacity and Grounds a Mathematical Model of Reversible T Cell Exhaustion

**DOI:** 10.64898/2026.05.05.722814

**Authors:** Aubrey Y. Liew, Ying Li, Joanina K. Gicobi, Jake B. Hirdler, Emilia R. Dellacecca, Haidong Dong

## Abstract

A central paradox in cancer immunotherapy is that patients harboring apparently exhausted or dysfunctional CD8^+^ T cells can exhibit rapid and durable responses to immune checkpoint inhibition (ICI). Although such rebound responses do not occur in all ICI-treated patients, resolving this paradox is essential for identifying and therapeutically maximizing reversible T cell exhaustion before clinical benefit is achieved. Here, we identify malic enzyme 1 (ME1) as a key molecular determinant of a latent, epigenetically poised effector CD8^+^ T cell state that is both necessary and sufficient for responsiveness to ICI. Genetic loss of ME1 abolishes therapeutic efficacy despite intact checkpoint blockade, whereas enforced ME1 expression enables robust antitumor responses even in otherwise ICI-resistant tumor models. Chromatin accessibility profiling, together with bulk and single-cell transcriptomic analyses, demonstrates that ME1 preserves effector readiness by maintaining latent effector capacity as a measurable biological state. These findings also experimentally ground a mechanistically interpretable dynamical model of T cell exhaustion, in which latent effector capacity, E(t), evolves according to a first-order differential equation shaped by activation history and decay. By defining latent effector capacity as a quantifiable, history-dependent state variable, our findings further enable the rational design of AI-driven models to predict patient-specific responsiveness and optimize therapeutic strategies in T cell–based immunotherapy. Together, our results redefine exhaustion as a state of masked potential governed by history-dependent latent effector capacity and provide a unified framework explaining rapid functional rebound, therapeutic heterogeneity, and the boundary between reversible and irreversible T cell exhaustion.

## INTRODUCTION

Most patients with advanced cancer harbor exhausted CD8^+^ T cells that are metabolically and epigenetically suppressed within the tumor microenvironment (TME) ^1-3^. These cells are characterized by diminished proliferative potential, impaired cytotoxic function, and sustained expression of inhibitory receptors such as PD-1. For many years, exhausted T cells were therefore viewed as terminally dysfunctional, i.e. locked into an irreversible differentiation state that precluded meaningful contributions to antitumor immunity. However, the clinical success of immune checkpoint inhibition (ICI), particularly therapies targeting the PD-1 pathway, has fundamentally challenged this view. A subset of patients mounts rapid and durable responses to ICIs^4-7^, in which T cells that appear functionally inert before treatment swiftly regain effector activity, including cytotoxicity against tumor cells, following PD-1 blockade^4,8,9^. This clinically observed paradox is difficult to reconcile with models in which exhaustion represents a complete loss of effector identity^4-6^. Although such rebound T cell responses do not occur in all ICI-treated patients, resolving this paradox is essential for identifying and therapeutically harnessing reversible T cell exhaustion, with the goal of improving the efficacy of ICI and other T cell–based immunotherapies.

Although exhausted CD8^+^ T cells have been associated with an epigenetic landscape indicative of terminal differentiation into a dysfunctional state^10-12^, recent studies demonstrate that, across the spectrum of exhausted CD8^+^ T cells, accessible chromatin is frequently retained at canonical effector loci^13-15^. These loci include genes encoding cytotoxic molecules and inflammatory cytokines^16,17^. Similarly, lineage-defining transcription factors associated with effector identity can remain bound to regulatory elements despite markedly attenuated downstream gene expression^18-20^. Collectively, these findings suggest that T cell exhaustion involves a dissociation between molecular competence and functional execution^15,21^, wherein the regulatory infrastructure required for effector function persists in a latent or masked state.

These observations motivate a conceptual distinction between two components of T cell state. The first is *latent effector capacity*, reflecting epigenetic accessibility and poised transcription factor networks accumulated through prior antigenic stimulation. This latent capacity evolves over relatively slow timescales and encodes the history-dependent imprint of past T cell priming and activation. Notably, this concept aligns with an early theoretical framework in cancer immunology described as “Invisibility of tumor immunity” ^22^, first articulated by Sir Frank Macfarlane Burnet, in which immune effector potential exists but is not phenotypically expressed^23,24^. The second component is *active effector output*, representing the realized transcriptional programs and functional execution, such as cytokine production and cytotoxicity, at a given moment in time. Active effector output is highly dynamic and can be readily assessed ex vivo in T cells isolated from patients; however, in vivo it may be masked by internal inhibitory signals, originally postulated by the Hellströms^25^ and molecularly defined as the PD-1/PD-L1 immune checkpoint pathway^26-29^.

While previous models of T-cell exhaustion have focused on differentiation trajectories or terminal fate decisions^30-33^, few have explicitly represented latent and history-dependent state variables alongside reversible exhaustion mechanisms. Without such a representation, it remains difficult to explain how PD-1 blockade can induce rapid functional rebound in some exhausted T cells yet fail completely in others, or to define the conditions under which exhaustion becomes irreversible. Addressing these questions requires a dynamical framework in which preserved effector competence (or capacity) and inhibitory masking are treated as distinct and interacting processes. In parallel, identifying molecular determinants of latent effector capacity is essential for experimentally grounding such models. Recent work has highlighted metabolic and epigenetic regulators that contribute to T-cell resilience under large tumor burdens^4,8,9^. Among these, malic enzyme 1 (ME1) has emerged as a candidate regulator associated with highly functional, therapy-responsive CD8^+^ T-cell states^4^. ME1 has been linked to enhanced metabolic fitness, effector readiness, and improved responses to cancer immunotherapy, suggesting that it may play a role in preserving the internal cellular programs required for effective immune responses.

In this study, we integrate experimental and theoretical approaches to resolve the paradox of reversible T cell exhaustion. Building on a mathematical framework that explicitly decouples latent effector capacity from active effector output, and models PD-1 signaling as a reversible masking mechanism acting on effector realization^34^, we experimentally ground this framework by demonstrating that ME1 defines and maintains a latent, epigenetically poised CD8^+^ T cell state that is both necessary and sufficient for responsiveness to ICI therapy. Using complementary genetic loss- and gain-of-function models of ME1 in CD8^+^ T cells, together with bulk and single-cell transcriptomic and epigenomic analyses, we find that ME1 preserves effector readiness by maintaining latent effector capacity as a measurable biological state. Our findings further anchor a mechanistically interpretable dynamical model of T cell exhaustion with the role of ME1 in T cells, in which latent effector capacity, denoted E(t), evolves according to a first-order differential equation, dE/dt, governed by activation history and decay. By directly linking an interpretable mathematical model to experimentally defined molecular states, this work provides a unified explanation for rapid functional rebound, heterogeneous therapeutic responses, and the emergence of irreversibility in T cell exhaustion.

## RESULTS

### ME1 establishes an epigenetically poised and effector-ready CD8^+^ T Cell state

ME1 (malic enzyme 1) is an NADP^+^-dependent enzyme that catalyzes the conversion of malate to pyruvate while generating NADPH^35,36^, a central reducing currency that sustains cellular antioxidant systems and redox homeostasis. From there, ME1 plays a well-established role in maintaining cellular redox homeostasis and protecting against oxidative stress^35,37^. However, its function in CD8^+^ T cells, particularly within the oxidative and nutrient-restricted TME, has remained largely unexplored. To determine whether ME1 is utilized by tumor-reactive CD8^+^ T cells in vivo, we assessed endogenous ME1 expression in CD8^+^ tumor-infiltrating lymphocytes (TILs). CD8^+^ TILs were isolated from MC38 tumors (a mouse colon cancer) at day 12 following tumor inoculation, a time point previously shown to coincide with peak effector T-cell accumulation^38,39^. Our flow-cytometric analysis of intracellular levels of ME1 revealed both an increased frequency of ME1^+^ CD8^+^ T cells and elevated ME1 protein abundance (mean fluorescence intensity, MFI) in CD8^+^ TILs compared with splenic CD8^+^ T cells from the same mice (**Fig. 1A–D**), suggesting ME1 upregulation in CD8^+^ T cells is associated with their presence in the TME, where they may have been activated through tumor antigen stimulation. To determine whether ME1 upregulation in CD8^+^ T cells is directly induced by T-cell activation, splenic CD8^+^ T cells were stimulated in vitro with anti-CD3/CD28 antibodies for up to 48 hours. As expected, T cell activation progressively increased ME1 expression, rising from a basal level to markedly higher expression states (**Fig. 1E–G**). Together, these results demonstrate that ME1 expression is activation-inducible and suggest that ME1 upregulation represents an adaptive feature of CD8^+^ T cells within the TME.

**Fig. 1.**
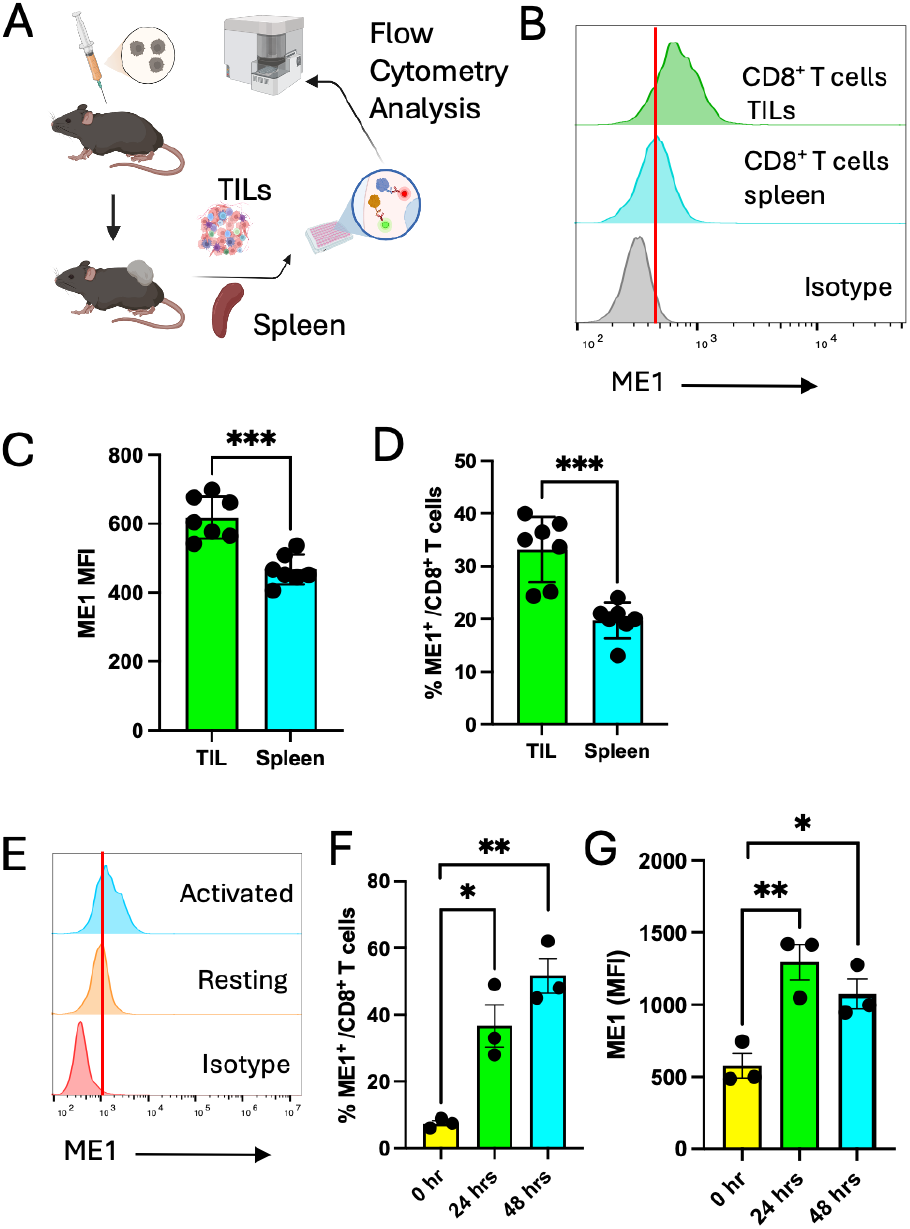
ME1 expression is induced upon CD8^+^ T-cell activation in peripheral and tumor compartments. (A) Schematic of ME1 expression assay. (B) Representative histogram of flow cytometry analysis of ME1 expression in CD8^+^ T cells isolated from tumor-infiltrating lymphocytes (TILs) and spleen on day 12 after MC38 tumor injection. (C-D) A summary of MFI (mean fluorescence intensity) of ME1 expression (C) or % ME1^+^ (D) in CD8^+^ T cells (n=7 mice). (E–G) ME1 expression in splenic CD8^+^ T cells following ex vivo activation with anti-CD3/CD28 antibodies at indicated times. (E) Representative flow cytometry histograms demonstrate upregulation of ME1 upon activation. (F-G) A summary of % ME1^+^ (F) or MFI (G) in CD8^+^ T cells (n=3 mice). For quantified comparisons, samples were analyzed using a two-tailed unpaired *Student’s t* test. Data are presented as mean ± SEM from two independent biological experiments. * P<0.05, ** P<0.01, *** P<0.001.

ME1 lies at the interface of mitochondrial TCA cycle metabolism and cytosolic NADPH production^37,40,41^, suggesting a potential role in coordinating redox balance and anabolic pathways that support T cell activation. Metabolites associated with ME1 activity, including pyruvate-derived acetyl-CoA, have been implicated in histone acetylation^42-44^, a chromatin modification linked to transcriptional accessibility. However, it is not clear whether and how ME1 contributes to epigenetic regulation of effector gene expression in CD8^+^ T cells. To address this possibility, we examined chromatin accessibility and transcriptional profile in CD8^+^ T cells by performing ATAC-seq and RNA-seq on cells isolated from the spleens of CD8-specific ME1 transgenic mice and littermate controls.

To examine how ME1 programs latent effector capacity in CD8^+^ T cells prior to tumor exposure, we isolated splenic CD8^+^ T cells from naïve mice and transiently activated them ex vivo with anti-CD3 and anti-CD28 antibodies for 24 hours, followed by bulk ATAC-seq analysis. Volcano plot analysis revealed numerous genomic loci with significantly increased chromatin accessibility in ME1-overexpressing CD8^+^ T cells, including prominent chromatin-open regions (CORs) associated with canonical cytotoxic effector genes such as *Gzmb, Ifng, Prf1*, and *Nkg7* (**Fig. 2A**). Consistent with this effector-biased chromatin landscape, CORs were increased at loci encoding transcription factors required for effector differentiation and lineage fidelity (*Klf2*)^45,46^, costimulatory signaling (*Cd28*), and memory cell maintenance (*Foxo1*)^47-49^, while accessibility at the *Ets1* locus, a transcriptional regulator known to suppress cytotoxic effector programs including *Nkg7*^*50*^, was reduced in CD8-ME1 transgenic T cells compared with controls (**Fig. 2A**). Notably, CORs at *Tox* and *Tcf7*, transcriptional regulators with opposing roles in T cell exhaustion^20^ and stem-like maintenance^51^, were also increased in ME1-overexpressing CD8^+^ T cells.

**Fig. 2.**
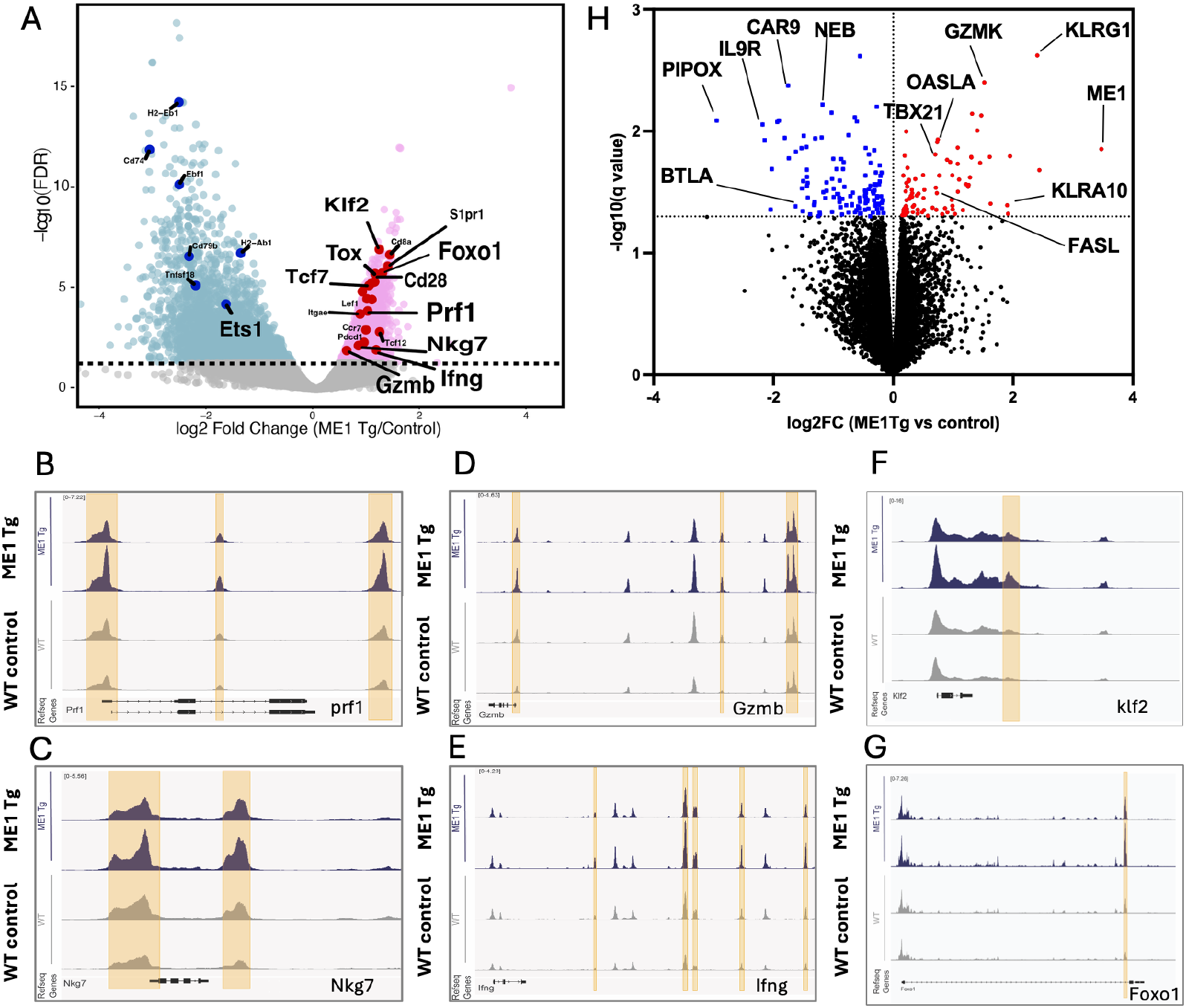
ME1 overexpression primes effector loci in CD8^+^ T cells. (A) Volcano plot showing gene-level differential chromatin accessibility in activated CD8^+^ T cells from ME1 transgenic (ME1 Tg) versus control mice. The x-axis indicates log_2_ fold change (ME1 Tg/control), and the y-axis denotes –log_10_ (adjusted *P* value) (n = 3 mice per group). Genes with an adjusted P value ≤ 0.05 are highlighted in pink and genes of interest in red. (B–G) Representative open chromatin region (OCR) tracks corresponding to selected effector-associated loci from panel (A), visualized in activated CD8^+^ T cells isolated from ME1 Tg (blue, top) or control (gray, bottom) mice using the Integrative Genomics Viewer (n = 2 mice per group). (H) Bulk RNA-sequencing analysis of splenic CD8^+^ T cells isolated from ME1 Tg and control mice following brief ex vivo activation. Statistically significant upregulated genes (red) and downregulated genes (blue) marked at significant cutoff of -log10(p-value<0.05) ≥ 1.3, dotted lines).

Closer inspection of representative ATAC-seq tracks demonstrated that ME1 overexpression broadly enhanced chromatin accessibility across multiple regulatory elements flanking hallmark cytotoxic effector genes. Specifically, ME1-overexpressing CD8^+^ T cells exhibited marked accessibility gains at promoter-proximal and intronic regulatory regions of *Prf1, Nkg7*, and *Foxo1* (**Fig. 2B–C, G**). Similar accessibility increases were observed at *Gzmb, Ifng*, and *Tfl2*, where ME1 overexpression induced distinct, sharply defined peaks across enhancer regions highlighted in tan in the genome browser views (**Fig. 2D–E, F**). Collectively, these data indicate that enforced ME1 expression broadly preserves chromatin accessibility across multiple T cell differentiation programs while selectively biasing the regulatory landscape toward effector lineage competence.

Consistent with these ATAC-seq findings, bulk RNA-seq analysis revealed that ME1 transgenic CD8^+^ T cells exhibited a poised effector transcriptional program, characterized by increased expression of *Tbx21* (T-bet), a master regulator of effector differentiation; effector-priming molecules such as *Gzmk* and *Fasl*; and *Oas1a*, a type I interferon–stimulated gene associated with interferon-driven immune priming (**Fig. 2H**). Together, these findings suggest that ME1 acts through epigenetic and transcriptional mechanisms to establish a state of latent effector capacity in CD8^+^ T cells, thereby promoting effector output upon subsequent activation.

### ME1-dependent latent effector capacity is functionally convertible into tumor control

Building on the poised effector capacity established by ME1 in splenic CD8^+^ T cells, we next examined how ME1 programs CD8^+^ T cell effector potential within the TME. To this end, we performed single-cell RNA-sequencing (scRNA-seq) analysis of CD8^+^ TILs isolated from MC38 tumors at a time point corresponding to peak effector cell accumulation. Notably, TILs from ME1 transgenic (Tg) mice exhibited a pronounced expansion of cluster 2, an effector-associated population, compared with TILs from control mice (**Fig. 3A**, red circles). Dot-plot analysis demonstrated that cluster 2 was enriched for high expression of key cytotoxic effector genes (*Gzmb, Prf1, Nkg7*), together with exhaustion-associated markers (*Pdcd1, Lag3, Tox*) (**Fig. 3B**), a pattern further supported by feature-plot visualization (**Fig. 3C**). This transcriptional profile of CD8^+^ TILs closely mirrored their epigenomic landscape of splenic activated CD8^+^ T cells, as ATAC-seq analysis revealed increased chromatin accessibility at loci encoding both cytolytic machinery and exhaustion markers in ME1 Tg mice compared with controls. Together, these findings indicate that ME1 upregulation promotes a sustained, high-potency effector program even within exhaustion-prone T cells in the TME.

**Fig. 3.**
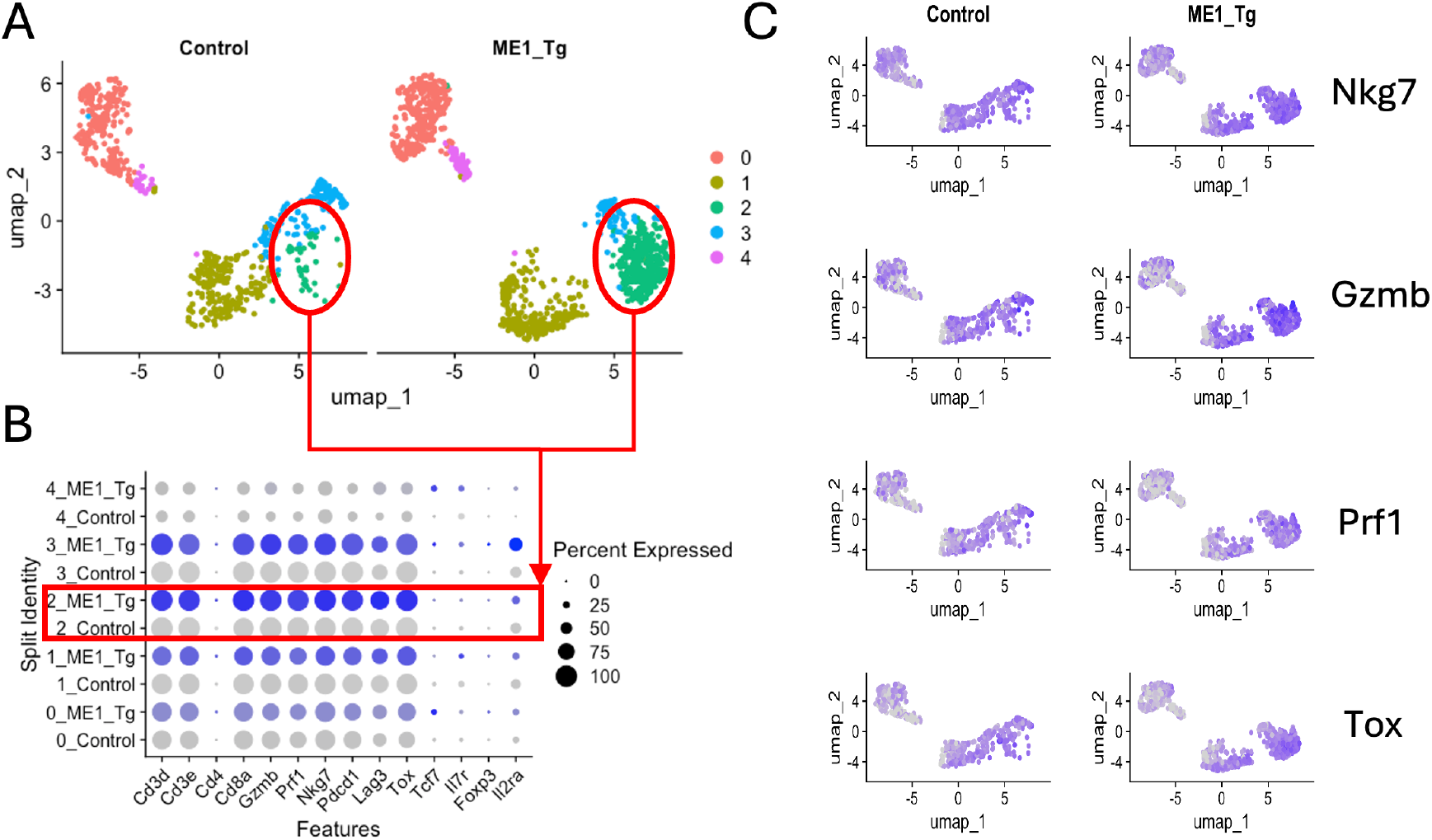
ME1 promotes an enhanced effector program in tumor-infiltrating CD8^+^ T cells. (A) UMAP projections of tumor-infiltrating CD8^+^ T cells from MC38-bearing control mice (left) and ME1 transgenic (ME1 Tg) mice (right), showing five transcriptionally distinct clusters (0–4). (B) Dot plot summarizing expression of selected genes across the five clusters. Dot color reflects average gene expression (blue, ME1 Tg; gray, control), with darker shading indicating higher expression levels. Dot size represents the percentage of cells within each cluster expressing the indicated gene. (C) Feature plots showing UMAP distributions and expression intensities of representative cytotoxic effector markers in CD8^+^ tumor-infiltrating lymphocytes from control (left) and ME1 Tg (right) mice. Gene expression levels are shown on a gradient scale, with darker blue indicating higher expression.

Given that CD8^+^ T cells resemble a state of latent effector capacity charged by ME1 upregulation, we next determined whether these pre-charged effecter capacity will give rise to potent cytotoxic TILs capable of superior tumor control. To test that, we evaluated MC38 tumor growth, an immunogenic colon cancer model, in CD8-specific ME1 transgenic (Tg) mice and littermate controls. As expected, the tumor growth was markedly delayed, and tumor volumes were significantly reduced in CD8-ME1 Tg mice compared with controls (**Fig. 4A–B**), indicating that ME1-programmed CD8^+^ T cells confer augmented antitumor activity.

**Fig. 4.**
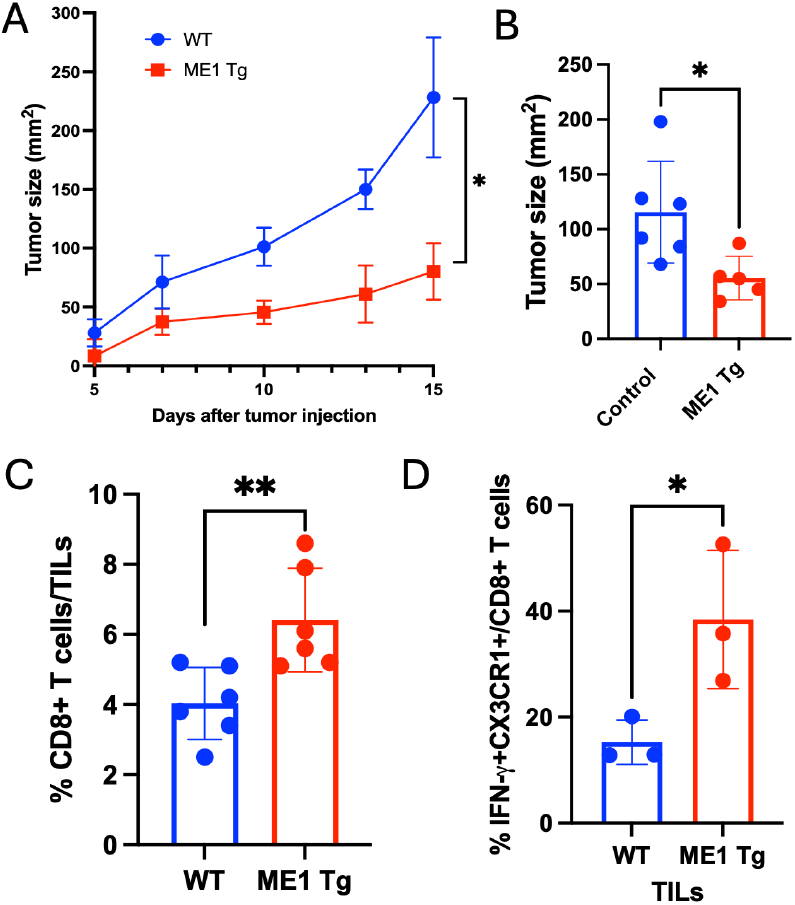
Antitumor immunity is enhanced in CD8^+^ T cell–specific ME1 transgenic mice. (A–B) MC38 tumor growth curves (A) and tumor sizes measured on day 15 (B). Statistical significance was assessed using two-way ANOVA for tumor growth (A, n = 5) and an unpaired two-tailed Student’s *t* test for tumor size (B, n = 5–6). (C–D) Frequency of CD8^+^ T cells among tumor-infiltrating lymphocytes (TILs) (C) and the proportion of IFN-γ^+^CX3CR1^+^ CD8^+^ T cells in TILs (D) on day 12 after tumor injection. Statistical analysis was performed using an unpaired two-tailed Student’s *t* test (n = 3–5). *P < 0.05; **P < 0.01.

This increased tumor control correlated with a significant expansion of CD8^+^ T cells within the TME (**Fig. 4C**) and an elevated frequency of functional CD8^+^ TILs, defined by IFN-γ and CX3CR1 expression, in CD8-ME1 Tg mice at day 12 following tumor inoculation compared with control mice (**Fig. 4D**). These findings provide direct evidence that the epigenetically and transcriptionally poised effector state established by ME1 upregulation in CD8^+^ T cells can be effectively translated into enhanced effector function within the TME, thereby driving tumor control.

To further test whether the ME1-primed effector state confers a T cell–intrinsic advantage to CD8^+^ T cells, we performed adoptive transfer experiments. CD8^+^ TILs were isolated from CD8-specific ME1 transgenic mice and littermate controls bearing MC38 tumors (**Fig. 5A**). Following three weeks of ex vivo expansion with IL-2 and anti-CD3/CD28 stimulation, ME1 Tg TILs expanded more robustly than control TILs (**Fig. 5B**), suggesting that ME1 can also enhance CD8^+^ T cell survival and/or proliferative capacity.

**Fig. 5.**
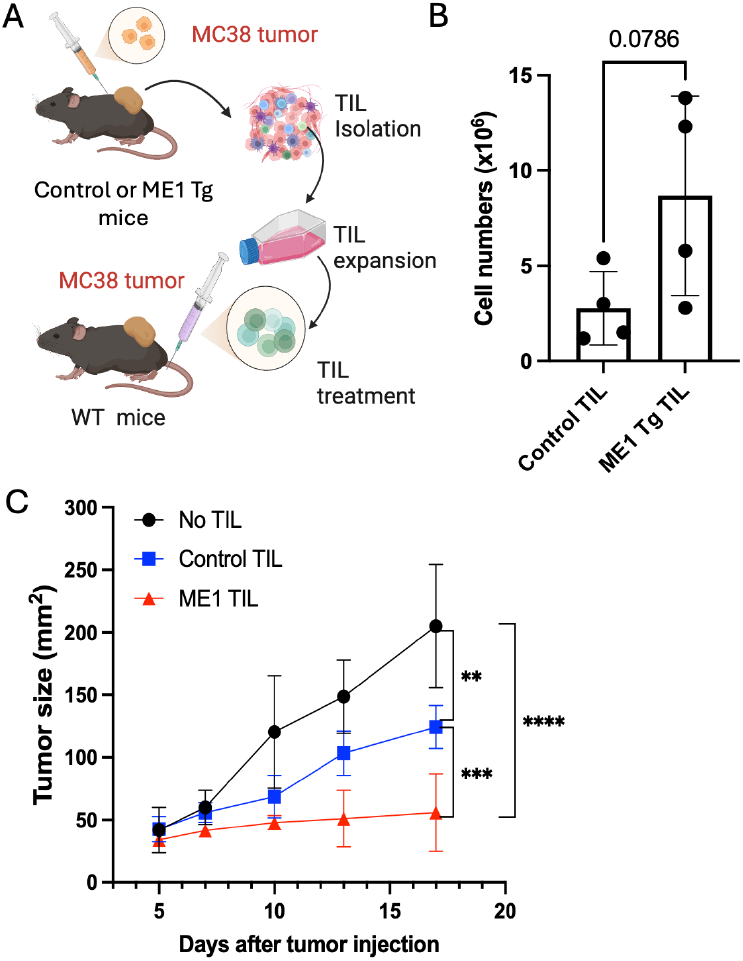
T cell–intrinsic ME1 upregulation is transportable in adoptive T cell transfer therapy. (A) Schematic diagram illustrating tumor-infiltrating lymphocyte (TIL) expansion and adoptive transfer therapy. (B) TILs isolated from tumors were expanded ex vivo for 3 weeks in the presence of IL-2 and anti-CD3/CD28 stimulation. Live cell numbers were quantified at the end of culture. Each dot represents pooled TILs from three mice within one experiment. Bar graphs summarize results from four independent experiments. Statistical significance was determined using an unpaired two-tailed Student’s *t* test. (C) MC38 tumor-bearing mice were treated with expanded TILs (1 × 10^5^ cells per mouse) via peritumoral injection on days 5 and 8 following tumor implantation. Tumor growth was analyzed by two-way ANOVA. **\*\***P < 0.01; **\*\*\***P < 0.001; **\*\*\*\***P < 0.0001 (*n* = 3–4 mice per group).

When these expanded TILs were adoptively transferred into secondary recipients with established MC38 tumors, ME1 Tg TILs conferred dramatic tumor control compared with control TILs (**Fig. 5C**). Together, these findings demonstrate that ME1 programs a stable, cell-intrinsic effector advantage in CD8^+^ T cells that is maintained through ex vivo expansion, transferable to a new host, and sufficient to drive enhanced antitumor activity.

### CD8^+^ T cell-associated ME1 is required and sufficient for effective ICI–mediated antitumor immunity

To test whether CD8^+^ T cell-associated ME1 in TME is required in responsiveness to immune checkpoint inhibition (ICI), we generated CD8^+^ T-cell–specific conditional ME1 knockout (CD8cre-ME1 fl/fl) mice on a C57BL/6 background. Using an ICI–responsive MC38 tumor model^52,53^, we evaluated the efficacy of PD-1 blockade in the absence of ME1 expression in CD8^+^ T cells (**Fig. 6A**). In the absence of immunotherapy, tumor growth was comparable between CD8^+^ T cell-specific ME1 conditional knockout (KO) mice and control littermates (**Fig. 6B**). In striking contrast, ICIs failed to confer therapeutic benefit in ME1 KO mice, whereas robust antitumor responses were observed in control mice receiving the same treatment (**Fig. 6C–D**). These findings demonstrate that ME1 expression in CD8^+^ T cells is required for effective ICI–mediated antitumor immunity and indicate that ICI alone is insufficient to overcome the loss of ME1-dependent, T-cell–intrinsic effector programs.

**Fig. 6.**
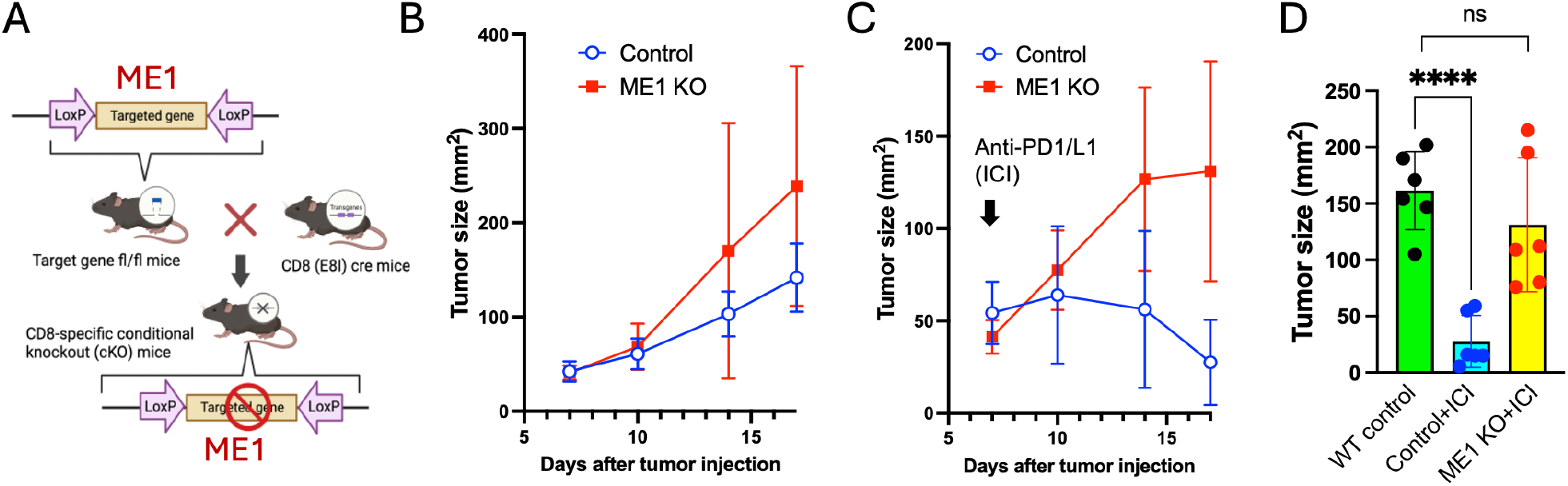
Loss of tumor response to immune checkpoint inhibition in CD8-specific ME1 knockout mice. (A) Schematic illustrating the generation of CD8-specific ME1 conditional knockout (CD8-ME1 KO) mice. (B–C) MC38 tumor growth in control (ME1 fl/fl) or ME1 KO mice without (B) or with immune checkpoint inhibitor (ICI) therapy (C, arrow). ICI therapy consisted of intraperitoneal injections of anti-PD-1 and anti-PD-L1 antibodies (G4/10B5 clones, Millipore Sigma, 100 µg each) administered every other day starting on day 6 after tumor implantation. Representative of two independent experiments is shown. (D) Summary of tumor size on day 18 (n = 6 mice per group). Data were analyzed using a two-tailed unpaired Student’s *t* test; ****P < 0.0001.

We next assessed whether enforced ME1 expression is sufficient to enable responsiveness to ICI therapy in resistant tumor contexts. To that end, we produced CD8^+^ T cell-specific ME1 transgenic mice (CD8-ME1 Tg mice) by breeding our pCAG2-ME1-ROSA26-IRES-tdTomato mice with CD8 (E8I) cre mice (**Fig. 7A**). In this model, the Cre deletes the stop codon before ME1 and allows ME1 to be constitutively expressed at the ROSA26 site that is marked by tdTomato (**Fig. 7B**), which can be used as a proxy marker to identify ME1 overexpression in CD8^+^ T cells. Since B16F10 tumor is poorly immunogenic and does not respond to ICI therapy ^7,54,55^, we used this model to test whether ME1 upregulation produces a stronger CD8^+^ T cell response to control B16F10 tumors upon ICI therapy.

**Fig. 7.**
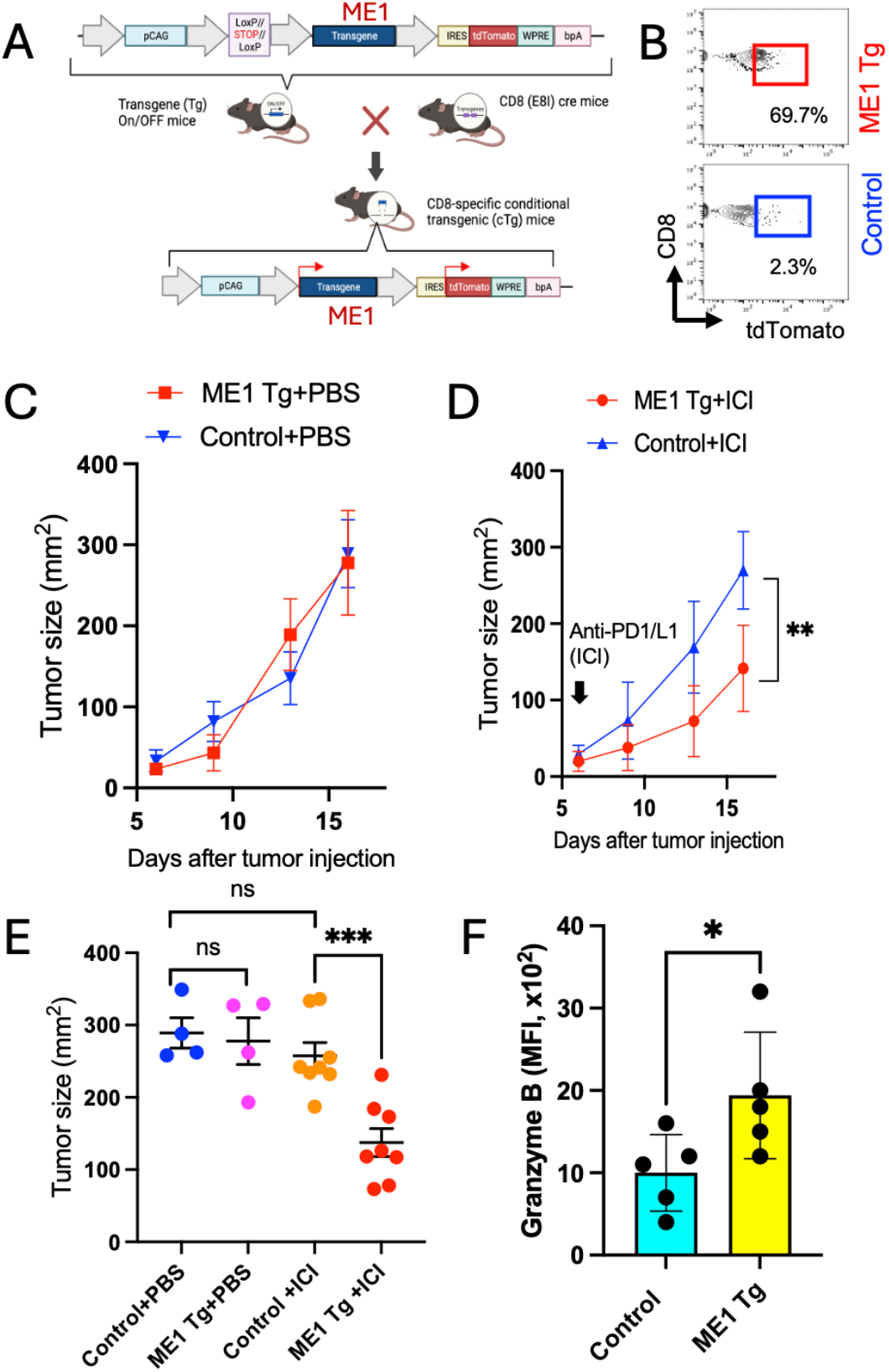
ME1 upregulation enhances the efficacy of immune checkpoint inhibition. (A) Schematic illustrating the generation of CD8-specific ME1 transgenic (ME1 Tg) mice. (B) Representative flow cytometry showing tdTomato expression as a surrogate for ME1 overexpression in CD8^+^ tumor-infiltrating lymphocytes (TILs). (C–D) Average growth curves of B16F10 tumors in control and CD8-ME1 Tg mice treated with PBS (C) or combined anti-PD-1/anti-PD-L1 antibodies (100 μg each per dose) administered every other day starting on day 6 (arrow) after tumor implantation. Tumor growth was analyzed by two-way ANOVA (D, n = 6, **P < 0.01). (E) Tumor sizes were measured at endpoint. Data were analyzed using an unpaired two-tailed *t* test (n = 4–8, ***P < 0.001). One of two independent experiments is shown. (F) Flow cytometric analysis of granzyme B (GZMB) protein expression in CD8^+^ TILs cells from B16F10 tumors in ME1 Tg and control mice on day 12. Data were analyzed using an unpaired two-tailed Student’s *t* test (*\** P< 0.05; n = 5 mice per group).

Although the overall growth of B16F10 tumors were comparable between CD8-ME1 Tg mice and control mice treated with PBS (vehicle control) (**Fig. 7C**), the tumor growth of B16F10 was suppressed and the tumor sizes decreased upon ICI therapy in CD8-ME1 Tg mice compared to control mice (ME1-ROSA26 without CD8 cre) (**Fig. 7D-E**). Accordingly, the expression of cytotoxic effector molecule granzyme B also increased in the ICI-treated CD8^+^ TILs of ME1 Tg mice compared to control mice (**Fig. 7F**). This result suggests that ME1 upregulation leads to a stronger antitumor activity of CD8^+^ T cells in responses to ICI therapy. This gain-of-function phenotype demonstrates that ME1 enhances a responsive and effective cytotoxic CD8^+^ T-cell state capable of supporting therapeutic efficacy through elimination of resistant tumor cells.

### Experimental grounding of a mechanistically interpretable model of T cell exhaustion

To interpret these findings within a quantitative framework, we build on our previously established model describing the dynamic progression of T cell exhaustion and the window of functional reversibility ^34^. In this model, latent effector capacity, *E*(*t*), evolves according to a minimal first-order differential equation:

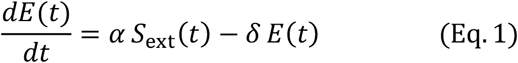

where *S*_ext_(*t*) captures combined external activation inputs, including antigen stimulation, co-stimulation and cytokine signals. The parameter *α* reflects the efficiency with which these signals are converted into durable, epigenetically encoded effector capacity, whereas *δ* represents the slow decay of this capacity associated with exhaustion. Thus, *E*(*t*) integrates the cumulative history of T cell activation, co-stimulation, and inflammatory signaling, providing a unifying quantitative link between observed molecular priming and long-term functional potential.

From the pattern of ME1 expression in a highly immunogenic MC38 tumor model, we demonstrate a strong association between tumor antigen–driven TCR signaling and ME1 upregulation in CD8^+^ tumor-infiltrating lymphocytes (TILs) (**Fig. 1B–D**). This relationship is further validated by ex vivo TCR stimulation assays (**Fig. 1E–G**). Together, these data indicate that ME1 upregulation in CD8^+^ T cells is tightly coupled to TCR stimulation. In this context, ME1 may function to transduce TCR signaling into a metabolic and epigenetic state that “charges” latent effector capacity. Complementary evidence is provided by ME1 transgenic (Tg) mice, in which constitutively elevated ME1 expression primes latent effector potential through chromatin remodeling and transcriptional reprogramming. This primed state is supported experimentally by integrated ATAC-seq and RNA-seq analyses of splenic CD8^+^ T cells (**Fig. 2**). Importantly, this priming is maintained within the TME, conferring poised effector readiness in situ, as demonstrated by single-cell RNA-seq analyses of CD8^+^ TILs (**Fig. 3**).

It is likely that ME1 functions primarily by increasing the parameter α, thereby accelerating the rate at which latent effector capacity, E(t), is “charged.” Once ME1 expression is upregulated or reinforced, it counteracts the decay term δE(t), shifting the system into a regime in which αS_ext_(t) ≫ δE(t). Under these conditions, E(t) accumulates progressively over time and reaches elevated levels within tumor tissues, where enhanced antitumor immunity can be established, which leads to a spontaneous delay in MC38 tumor growth relative to control mice, consistent with our experimental observations (**Fig. 4**). Notably, this “pre-charged” effector capacity within CD8^+^ TILs is transferable: adoptive transfer of TILs from ME1 Tg mice into secondary tumor-bearing hosts confers significantly greater antitumor activity compared with control TILs (**Fig. 5**).

The dynamic evolution of latent effector capacity, E(t), can therefore be intuitively represented by a water-tank model (**Fig. 8**). In this analogy, the input term αS_ext_(t) functions as a hose supplying water to the tank, while the decay term δE(t) represents a leak at the bottom. The total water volume corresponds to E(t), which is dynamically determined by the balance between inflow and outflow. Within this framework, ME1 is proposed to enhance effector capacity indirectly by increasing *α*, which amplifies the efficiency of converting external stimulation into effector capacity accumulation. In this model (**Fig. 8B**), *α* represents the size of inflow hose or equivalently, the rate of inflow through the hose.

**Fig. 8.**
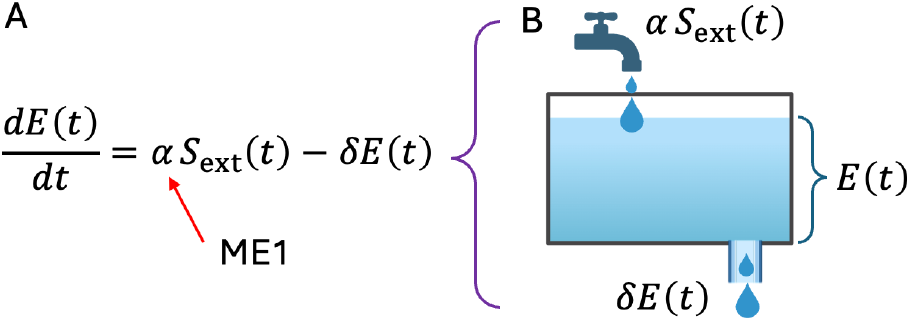
Schematic water-tank model illustrating Equation 2. (A) Latent effector capacity *E*(*t*) is determined by the balance between external stimulation and intrinsic decay. (B) In the water-tank analogy, external stimulation *S*_ext_(*t*)(depicted as a water hose) is converted into effector accumulation through the scaling parameter *α*. Effector capacity decays proportionally to its current level at rate *δE*(*t*), represented as a leak (hole) at the bottom of the tank. The resulting dynamics of *E*(*t*)(represented by the water volume) are therefore governed by the balance between stimulus-dependent inflow *αS*_ext_(*t*) and leakage *δE*(*t*). ME1 is indicated by increasing *α*.

To model immune checkpoint regulation, we represent PD-1–mediated inhibition using a Hill-type masking function:

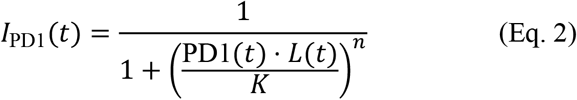

Here, *I*_PD1_(*t*) denotes the effective inhibitory input experienced by a T cell at time *t*, reflecting the strength of PD-1 checkpoint signaling in suppressing effector function. PD1(*t*) represents PD-1 receptor expression, *L*(*t*) denotes ligand availability (e.g., PD-L1), *K* defines the effective inhibitory threshold, and the Hill coefficient *n* parameterizes the cooperativity of PD-1–mediated suppression.

We next define realized effector activity, *A*(*t*), as the product of intrinsic latent effector capacity *E*(*t*) and checkpoint modulation via *I*_PD1_(*t*). This interaction is formalized by a multiplicative gating relationship:

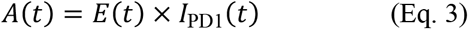

This formulation captures the principle that PD-1 signaling suppresses the *execution* of effector programs without directly altering the underlying latent effector capacity-namely, the epigenetic and transcriptional competence encoded in *E*(*t*). Within this framework, effective PD-1 blockade corresponds to removal of the inhibitory mask (PD1(*t*) → 0 and thus *I*_PD1_(*t*) → 1). In the limiting case of complete checkpoint blockade or absence of PD-1 signaling,

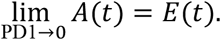

This limiting behavior explains the rapid functional rebound observed experimentally following PD-1 blockade in immune checkpoint inhibitor (ICI)–responsive tumor models such as MC38 (**Fig. 6C–D**), a process that may not require de novo epigenetic reprogramming at the time of therapy.

In contrast, in ME1-deficient CD8^+^ T cells, our experimental data indicate a collapse of the latent effector capacity variable *E*(*t*). Consequently, even when PD-1 signaling is pharmacologically blocked, effector output remains negligible:

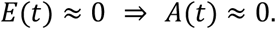

This result explains the failure of PD-1 blockade in ME1-deficient mice, even in the context of an ICI-responsive tumor model (MC38) (**Fig. 6C–D**) and supports a central assumption of our model: ICI unmasks pre-existing effector potential rather than generating it de novo. If PD-1 blockade could create effector potential *de novo*, ICI therapy alone would be sufficient to induce MC38 tumor regression even in the absence of ME1. Our data do not support this scenario.

Our experimental findings indicate that basal ME1 expression establishes and maintains a latent effector capacity *E*(*t*), whereas PD-1 signaling functions as a reversible inhibitory (“masking”) term *I*_PD1_(*t*). Observed effector activity therefore arises from the multiplicative interaction between *E*(*t*) and this reversible inhibitory state. Loss of ME1 collapses latent effector capacity, rendering ICI ineffective (**Fig. 6C–D**), whereas ME1 overexpression expands the domain of reversibility and enables therapeutic responses to checkpoint blockade even in otherwise resistant tumor contexts, such as B16F10 melanoma (**Fig. 7D–E**). Of note, in the B16F10 melanoma model, ME1 overexpression in CD8^+^ T cells, in the absence of ICI treatment, was insufficient to induce tumor regression (**Fig. 7C**). This finding indicates that enhanced ME1-dependent metabolic and epigenetic reprogramming alone cannot compensate for weak antigen-specific TCR signaling, which is characteristic of the low immunogenicity of B16F10 tumors, to initiate full effector differentiation and cytotoxic activity. Rather, ME1 overexpression appears to potentiate, but not autonomously trigger, CD8^+^ T cell effector function, underscoring its dependence on external activation cues (*S*_ext_(*t*)). From this perspective, ME1 functions to preserve chromatin accessibility and regulatory readiness at effector loci (**Fig. 2**), thereby establishing a molecular substrate for the latent effector capacity variable *E*(*t*).

Based on experimental data, ME1 is incorporated as an indirect regulator of effector capacity by modulating the efficiency parameter *α*. From there, we rewrite Eq. 1 in standard linear form,

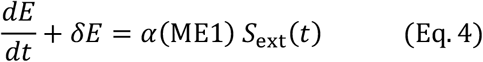

where *S*_ext_(*t*) denotes external stimulation, *δ* represents exhaustion-mediated loss, and *α*(ME1) captures the ME1-dependent efficiency with which stimulation is converted into latent effector capacity. Solving this equation using an integrating factor (*e*^*δt*^) yields the closed-form solution:

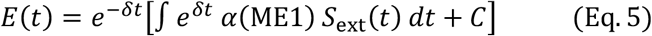

where *e*^*δt*^ represents the exhaustion discounting term and *C* denotes the initial effector capacity. This expression shows that *E*(*t*)is an exhaustion-discounted integral of prior stimulation, with ME1 entering as a multiplicative gain on the effective input to the latent effector capacity pool.

Since ME1 preserves chromatin accessibility and regulatory readiness at effector gene loci, it provides a molecular substrate for the latent effector capacity variable *E*(*t*). In the mathematical solution (Eq. 4-5), this role is reflected by the ME1-dependent scaling of *α*, which increases the amount of effector capacity accumulated per unit stimulation without altering the intrinsic decay rate *δ*. As a result, ME1 primarily influences latent, epigenetically encoded effector potential rather than instantaneous effector output. The model further predicts that ME1-dependent amplification of stimulation is required to maintain effective T cell responses under conditions of chronic or repetitive TCR engagement, such as within the TME or during chronic viral infection. Together, these findings provide direct experimental validation of *E*(*t*) as a biologically meaningful latent state variable and establish ME1 as a key metabolic determinant of reversible T cell effector competence.

## DISCUSSION

In this report, we address the T cell exhaustion paradox-namely, the rapid functional rebound of apparently dysfunctional T cells following immune checkpoint inhibitor (ICI) therapy in patients, by experimentally grounding a mathematical framework. As we previously reported^34^, this framework explicitly decouples latent effector capacity from active effector output, and here we identify ME1 as a key molecular determinant of this latent effector capacity.

Our data demonstrate that ME1 does not merely correlate with effector-like transcriptional signatures observed in clinical studies^4^, but instead programs a latent, epigenetically poised CD8^+^ T cell state that is both necessary and sufficient for responsiveness to ICI therapy. Loss- and gain-of-function studies reveal that ICI therapy fails in the absence of ME1 but becomes effective in otherwise non-responsive tumor contexts, such as the B16F10 model, when ME1 expression is enforced. Furthermore, ATAC-seq, bulk RNA-seq, and single-cell RNA-seq analyses establish that ME1 upregulation preserves chromatin accessibility and regulatory readiness at canonical effector loci upon TCR stimulation. These findings provide direct biological grounding for the latent effector capacity variable, E(t), in our model and support the interpretation of T cell exhaustion as a state of masked competence rather than irreversible loss of effector potential.

A key insight emerging from this framework is that immune checkpoint signaling functions primarily as a reversible gating or masking mechanism of effector realization, rather than as an immediate driver of epigenetic collapse or erasure ^11,33^. In the mathematical formulation, this principle is captured by a multiplicative interaction between latent effector capacity and PD-1–dependent inhibition. Our experimental data support this structure: ICI therapy unmasked robust effector function only when ME1-dependent latent effector capacity was preserved, whereas checkpoint blockade was ineffective once this capacity had collapsed. This distinction resolves a long-standing ambiguity regarding whether ICIs actively “reprogram” exhausted T cells or instead release pre-existing effector potential. Our results argue for the latter and formalize this concept in a quantitative, mechanistically interpretable manner through our mathematical model. Importantly, the mathematical model further predicts that T cell exhaustion is governed by history-dependent dynamics, with effective irreversibility emerging only after sustained inhibitory signaling destabilizes epigenetic self-maintenance ^10^. Although our study does not quantitatively define the precise bifurcation point separating reversible from irreversible exhaustion, both genetic perturbations and functional outcomes are consistent with the existence of distinct epigenetic regimes: ME1 loss abolishes responsiveness even under PD-1 blockade, whereas ME1 overexpression expands the domain of reversibility within otherwise resistant tumor microenvironments. Our findings suggest that therapeutic timing and preservation of latent effector capacity may be as critical as acute checkpoint inhibition itself.

Within this framework, the parameter *α* quantifies the efficiency with which external activation signals *S*_ext_ (*t*), including TCR stimulation, co-stimulation, and inflammatory cues, are converted into durable, epigenetically encoded effector capacity *E*(*t*). Biologically, *α* reflects the capacity of CD8^+^ T cells to translate transient activation signals into sustained chromatin accessibility, transcription factor occupancy, and regulatory readiness at effector loci. ME1 acts primarily along this axis: ME1 expression increases *α* by stabilizing chromatin accessibility and reinforcing transcriptional networks that encode effector competence, thereby enabling strong activation signals to efficiently accumulate a persistent latent effector state. In contrast, the parameter *δ* represents the slow, exhaustion-associated decay of latent effector capacity over time, capturing cumulative processes such as chronic antigen exposure, metabolic stress, and repressive chromatin remodeling. Importantly, *δ* operates on a slower timescale than acute effector execution and reflects structural erosion of epigenetic competence rather than immediate inhibitory signaling. In ME1-deficient CD8^+^ T cells, *δ* effectively dominates, leading to progressive collapse of *E*(*t*) even in the presence of ongoing antigenic stimulation (like in a high immunogenic tumor model of MC38). Collectively, the balance between *α* and *δ* determines whether latent effector capacity is maintained, expanded, or lost: when *αS*_ext_(*t*) > *δE*(*t*), *E*(*t*) accumulates, preserving a reversible exhaustion state that can be rapidly unleashed by checkpoint blockade; conversely, when *δ* dominates, *E*(*t*) decays toward zero, rendering ICIs ineffective. Thus, ME1 expands the domain of reversibility not by directly altering checkpoint masking, but by shifting the dynamical balance toward durable maintenance of latent effector capacity.

Placing these results in the broader context of T cell differentiation, our framework bridges concepts across exhaustion, memory formation, and immune resilience. Latent effector capacity shares defining features with stem-like or progenitor exhausted T cells ^56,57^, including preserved chromatin accessibility and responsiveness to ICIs, yet it is not reducible to surface phenotype or lineage designation. Instead, it represents an internal dynamical state established by the cumulative regulatory and epigenetic architecture acquired over time. By explicitly distinguishing this latent effector capacity from realized effector output, the model provides a unifying explanation for why exhausted T cells may appear inert in functional assays while retaining substantial rebound potential, and why this potential is ultimately finite. This separation clarifies both the mechanistic basis of therapeutic responsiveness and the limits of immune reinvigoration in chronic antigen settings.

Beyond its mechanistic insights, the framework established here provides a principled foundation for training artificial intelligence (AI) models to predict and design T cell–based immunotherapies for cancer. By experimentally grounding latent effector capacity *E*(*t*) as a biologically meaningful, slowly evolving state variable governed by a first-order dynamical equation (*dE*/*dt* = *αS*_ext_(*t*) − *δE*(*t*)), this study defines interpretable parameters that can be directly incorporated into machine-learning architectures. ME1 expression, chromatin accessibility, and transcriptional effector readiness can serve as measurable proxies for the loading efficiency *α* and decay rate *δ*, enabling AI models to distinguish reversible from irreversible exhaustion states rather than treating exhaustion as a static endpoint. Importantly, representing PD-1 signaling as a reversible masking function acting on, rather than redefining, *E*(*t*) allows predictive models to decouple therapies that unmask effector potential from those that must rebuild it. This separation reduces confounding in outcome prediction and imposes biologically grounded constraints that improve generalizability across tumor types and therapeutic modalities. More broadly, embedding experimentally validated state variables and dynamical rules into AI frameworks shifts immunotherapy modeling from correlation-based classification toward mechanistically supervised learning, enabling simulation of therapeutic timing, combination strategies, and patient-specific response trajectories. In this way, latent effector capacity emerges not only as a biological insight but also as a scalable intermediate representation for predictive and generative AI in cancer immunotherapy.

Several limitations of the present study merit discussion. First, while multiple datasets converge on ME1 as a determinant of latent effector capacity, the model does not claim that ME1 is the sole regulator of this state. Other chromatin regulators, transcription factors, and metabolic programs likely contribute and may define parallel or partially independent components of latent capacity. Second, the mathematical framework prioritizes interpretability and structural clarity over exhaustive molecular detail. Parameters governing epigenetic self-maintenance, decay, and bifurcation behavior are not directly measured here, and future longitudinal perturbation studies will be required to quantitatively calibrate these dynamics. Third, our analysis is cell-intrinsic and does not explicitly model population-level processes such as clonal replacement or selective expansion, which undoubtedly contribute to in vivo responses to ICI therapy. These processes are complementary rather than contradictory and incorporating them represents an important direction for multi-scale extension of the model.

In summary, this study establishes ME1 as a molecular determinant of latent effector capacity and provides experimental grounding for a mathematical framework that decouples epigenetic potential from inhibitory masking in exhausted CD8^+^ T cells. By resolving the apparent contradiction between preserved molecular competence and dysfunctional output, the framework offers a unified explanation for rapid rebound, therapeutic heterogeneity, and the point at which exhaustion becomes irreversible. More broadly, it illustrates how experimentally anchored dynamical models can transform descriptive immunology into predictive, mechanistically interpretable frameworks for cancer immunotherapy.

## METHODS AND MATERIALS

### Mice

All mouse experiments were carried out at Mayo Clinic in Rochester, Minnesota, and mice were maintained under specific pathogen-free conditions within the animal facility with all experimental protocols approved by the Institutional Animal Care and Use Committee (IACUC) at Mayo Clinic (IACUC: A00006353-21-R24). Purchased male and female 6-to 8-week-old C57BL/6 (B6) mice were allowed to acclimate to our housing facilities for at least one week. E8ICre mice (The Jackson Laboratory, C57BL/6-Tg(CD8a-cre)1Itan/J, strain #008766) with Cre recombinase under control of the CD8^+^ T cell specific E8I enhancer, were crossed to *ME1 loxP*-flanked mice (*ME1*^fl/fl^) and CAG2-NKG7-Rosa26-IRES-tdTomato mice (Rosa26-ME1), both were custom generated on a C57BL/6N background by Ingenious Targeting Laboratory, Ronkonkoma, NY) to induce deletion of *ME1* or overexpression of *ME1* in peripheral CD8^+^ T cells, generating a conditional knockout (KO) or transgenic (Tg) model. In the Tg mice, CD8 promotor driven Cre enzyme will remove stop codon within a new construct ahead of *ME1* gene in CD8^+^ T cells and enable bicistronic expression of *ME1* and tdTomato.

Upon recipient in our lab, future generations of mice were genotyped by DNA extraction from tail snips (DNeasy Blood and Tissue Kit, Qiagen, cat #69506) included in polymerase chain reaction (PCR) with GoTag Green Master Mix (Promega, catalog (cat) #M7123) and following protocol #35285 from The Jackson Laboratory for Cre-recombinase status. The *ME1*^fl/fl^ and Rosa26-ME1 status were confirmed by PCR on a SimpliAmp thermal cycler (Applied Biosystems™) and run on 1% TAE ReadyAgarose Precast gels with ethidium bromide (Bio-Rad, cat #1613050) and imaged on an iBright Imaging System (Invitrogen™). Homozygous dominate *ME1*-floxed mice, or homozygous expression of Rosa26-ME1, and heterozygous E8I-cre mice were included in experiments. Non-aged mice were utilized for experiments and mice were randomized to make average age as similar as possible between corresponding groups, while still also including both male and female mice in the same experiments, where possible. In tumor experiments mice were randomly assigned to treatment group by tumor size to normalize initial tumor sizes to the best of our abilities prior to treatment.

### Tumor model and treatment

MC38 (cat #SCC172) from Sigma Aldrich as validated cell lines. B16F10 murine melanoma cell line was purchased from ATCC (CRL-6475) and cultured using DMEM complete medium. All cell lines were maintained at 37°C with 95% relative humidity and 5% carbon dioxide and grown in Dulbecco’s Modified Eagle Medium (DMEM, Gibco, cat #11885-084). All media was made complete by addition of 10% fetal bovine serum (FBS, Gibco, cat #26400044, heat-inactivated in-house), 1X penicillin/streptomycin (Corning, cat #30-002-Cl), and 20 mM HEPES buffer (Corning, cat #25-060-CI). Mycoplasma detection of cell lines was performed using the PlasmoTest™ detection kit (InvivoGen, cat #rep-pt1) and authenticated by short tandem repeat profiling (B16 at American Type Culture Collection and MC38 at IDEXX). Cell lines were frozen in Recovery Cell Culture Freezing Medium (Gibco, cat #12648010) and stored for long-term storage in liquid nitrogen prior to thawing for experimental use. All adherent cells were lifted with 1X Trypsin-EDTA (Corning, cat #25-053-Cl) and neutralized with appropriate media prior to centrifugation. ICI therapy includes injection of anti-PD-1/PD-L1 (Millipore Sigma #MABC1132 (G4/PD-1) and #MABC 1131(10B5/PD-L1) at 100 ug of each through intraperitoneal injection every other day stating on day 6 after tumor injection.

### Murine cell isolation and activation

Upon humane euthanasia, as approved by the IACUC at Mayo Clinic, spleens were harvested from mice and homogenized using sterile glass slides and resuspended in RPMI 1640 and strained through a 70 mm filter (Corning, cat #431751) to obtain a single cell suspension. Red blood cells were lysed with ACK lysis buffer (made in the Preparation and Processing Lab (PPL), in the Department of Laboratory Medicine and Pathology, Mayo Clinic Rochester) prior to further isolations or analyses. Murine CD8^+^ T cells were isolated from splenocytes by EasySep™ negative immunomagnetic separation following manufacturer’s instructions (STEMCELL Technologies, cat #19853). Dissected tumors were homogenized with a Tumor Dissociation Kit (Miltenyi Biotec, cat #130-096-730) following manufacturer’s protocol and gentleMACS C Tubes (Miltenyi Biotec, cat #130-093-237) on a gentleMACS Octo Dissociator with Heater (Miltenyi Biotec), utilizing program 37C_m_TDK_1. Dissociated tumors were washed with RPMI 1640 and run through a 70 mm filter prior to downstream assays. TILs were isolated from tumor suspensions utilizing an EasySep™ magnetic CD45^+^ positive selection kit (STEMCELL Technologies, 100-0350) following manufacturer’s recommendations. Murine CD8^+^ T cells were activated with anti-CD3/CD28 Dynabeads (Gibco, cat #11452D) in complete RPMI 1640 following manufacturer recommendations. CD8^+^ T cells were seeded at 1×10^6^/mL in 24 well plates (Corning, cat #3526).

### RNA isolation

RNA was isolated from murine cells with TRIzol™ Reagent (Invitrogen™, cat # 15596026) following manufacturer’s recommendations. Chloroform (Thermo Scientific™, cat #326821000) was added to TRIzol resuspended samples prior to centrifugation at 12,000 x g for 15 minutes at 4ºC to obtain the aqueous layer. An equal volume of ethanol (Sigma-Aldrich, cat #459844) was added to the aqueous layer prior to inclusion in spin column-based RNA isolation following manufacturer’s protocol with addition of DNase (Zymo Research, cat #R1013). RNA quality and quantity was calculated by nanodrop (DeNovix DS-11) and stored at -80ºC prior to downstream assays including quantitative reverse transcription polymerase chain reaction (qRT-PCR) and bulk RNA-sequencing.

### Murine tumor infiltrating lymphocyte (TIL) transfer

CD8-specific ME1 Tg mice and control mice were subcutaneously injected with 1×10^6^ MC38 cells resuspended in sterile PBS into a shaved flank. Tumor measurements were performed biweekly with calipers and volume was calculated by an ellipsoid formula in which volume = length x width^2^ x 1/2. Mice were euthanized 12-days post tumor inoculation, in which tumors had become palpable and enough time for effector CD8^+^ T cell generation. Tumors were then dissociated, as described above. Dead cells were removed from single cell tumor digestion samples utilizing a dead cell removal kit (Miltenyi Biotec, cat #130-090-101), following manufacturer’s recommendations. Post-dead cell removal samples were then resuspended in complete RPMI 1640 at 1×10^6^ cells/mL and plated into 24-well culture plates with the addition of 6,000 IU/mL rhuIL2 and cultured for 72 hours at 37°C. TILs in culture were then stimulated and expanded with anti-CD3/CD28 beads (as above) following manufacturer recommendations and cultures were split upon confluency with fresh addition of activation beads and IL2. TILs were allowed to expand for 16 days post-culture prior to pooling within genotype and injection into recipient mice. Wildtype C57BL/6 mice were used as recipient mice and received 0.5×10^6^ MC38 cells resuspended in sterile PBS as a subcutaneous injection into a shaved flank. After seven days of growth, with palpable tumor measurements, mice were randomized into treatment groups for normalized tumor sizes across groups, and recipient mice received TILs from either CD8-specific ME1 Tg mice or control mice (post-activation bead removal by magnetic separation and two times wash with sterile PBS) at a dose of 1×10^6^ or PBS alone (no TIL treatment) by peri-tumoral injection. Mice received a second dose two days later at 9 days post-tumor inoculation. Tumor measurements were performed biweekly with calipers and volume was calculated by an ellipsoid formula in which volume = length x width^2^ x 1/2. Mice were euthanized upon reaching humane endpoint based off IACUC approved maximum tumor volume, ulceration, or weight-loss criteria.

### Spectral flow cytometric analysis

Single cell suspensions were washed in PBS prior to plating into a 96-well U-bottom plate at 1×10^6^ cells per stain. Cell viability is determined by staining with 1:1,000 Ghost Dye™ Violet 510 (Tonbo® sold by Cytek, cat #13-0870-T100) in sterile PBS following manufacturer’s recommendations. Cells were then washed twice with FACS buffer (PBS + 2% FBS + 2 mM EDTA (Invitrogen™, cat #15575020)) prior to resuspension in Fc block (TruStain FcX™ (anti-mouse CD16/32), BioLegend, cat #101320) following manufacturer’s recommendations. Surface antibodies were prepared in Brilliant Stain Buffer (BD Biosciences, cat #563794) and cells were resuspended in 100 µL of surface master mix for 30 minutes on ice. Cells were then washed 3 times with FACS buffer prior to fixation and permeabilization utilizing a FoxP3/Transcription Factor Staining Buffer kit (Invitrogen™, cat #00-5523-00) with 45-minute fixation on ice and permeabilization following kit recommendations. Intracellular antibodies were diluted in 1X permeabilization buffer and incubated on ice for 60 minutes. Cells were then washed 3 times with 1X permeabilization buffer and resuspended in 1% paraformaldehyde (PFA, Electron Microscopy Sciences, cat #15710) in PBS until samples were run on the cytometer the next day. Antibodies and their dilutions can be found in **Table 1**. All samples were run on a five laser Cytek Aurora spectral flow cytometer utilizing SpectroFlo® software. Quality control (QC) was run upon start-up of the Aurora each day utilizing SpectroFlo® QC beads (Cytek, cat #B7-10001). Spectral unmixing was calculated with unstained cells, viability-stained cells, and all other markers demonstrating positive signal on UltraComp eBeads Plus compensation beads (Invitrogen, cat #01-2222-42) prepared fresh for each experiment. Flow files were analyzed with FlowJo software (BD, v10.8.0). Gates were set on FSC-H vs SSC-H to determine overall cell population, followed by singlet inclusion by FSC-A vs FSC-H, then live cells by FSC-A vs Ghost Dye Violet 510 (gated on the negative), and CD8 vs TCR-beta double positive, prior to further gating. As panels were designed and optimized, FMOs of all surface markers were included and isotype controls for intracellular markers were included. Gates were adjusted as needed to capture mild inter-individual differences. Absolute cell counts were back calculated from total cell number as determined by trypan blue staining.

**Table 1.** Major reagents used in flow cytometry analysis.

| Reagent | Conjugate /Channel | Clone | Vendor | Catalog number | Dilutions |
| --- | --- | --- | --- | --- | --- |
| Ghost Dye V510 |  |  | Cytek Biosciences | SKU 13-0870-T100 | 1:1000 |
| Anti-mouse TCRβ | PerCP /Cyanine5.5 | H57-597 | Biolegend | 109227 | 1:200 |
| Anti-mouse CD8 | BUV395 | 53-6.7 | BD Biosciences | 569181 | 1:200 |
| Anti-mouse CX3CR1 | BV650 | SA011F11 | Biolegend | 149033 | 1:100 |
| Anti-Granzyme B | PE | QA18A28 | Biolegend | 396406 | 1:100 |
| Anti-IFN-gamma | BV480 | XMG1.2 | Thermo Scientific | 14-7311-81 | 1:100 |
| Anti-ME1 | PE | C-6 | Santa Cruz Biotechnology | sc-365891 | 1:100 |

### Bulk RNA and ATAC sequencing

All sequencing experiments were run in the Genome Analysis Core (GAC) within the Center for Individualized Medicine at Mayo Clinic Rochester. RNA was extracted in the lab, as described above, from 1×10^6^ nucleofected and activated (or rested) CD8^+^ T cells and the GAC measured total RNA concentration and QC by Qubit fluorometry (ThermoFisher Scientific) and the Agilent Fragment Analyzer. Based on the RNA quality, cDNA libraries were prepared using up to 250 ng of total RNA according to the manufacturer’s instructions for the Illumina TruSeq Stranded mRNA Prep (Illumina). The concentration and size distribution of the completed libraries were determined with the Agilent TapeStation’s D1000 ScreenTape and Qubit fluorometry. Libraries were normalized to 2 nM and sequenced following Illumina’s standard protocol for the NovaSeq 6000. The libraries were loaded across a SP flow cell at a final concentration of 400 pM. The flow cell was sequenced as 100 × 2 paired end reads using the NovaSeq SP Reagent Kit v1.5 and NovaSeq Control Software v1.8.0. Base-calling was performing using Illumina’s RTA version 3.4.4. For mouse samples, cDNA libraries were prepared using up to 100 ng of total RNA according to the manufacturer’s instructions for the Illumina Stranded mRNA Prep, Ligation (Illumina). The concentration and size distribution of the completed libraries were determined with the Agilent TapeStation’s D1000 ScreenTape and Qubit fluorometry. The libraries were normalized to 2 nM and sequenced following Illumina’s standard protocol for the NovaSeq X Series. The libraries were pooled and loaded onto a lane of a 10B flow cell at a final concentration of 60 pM. Following Illumina’s standard protocol, the flow cell was sequenced with 100 × 2 paired end reads using the NovaSeq X Plus Control Software v1.3.0 and RTA4.

For ATAC-seq, CD8^+^ T cells were collected in low-binding microcentrifuge tubes (USA SCIENTIFIC, cat #4043-1081) and washed in ice-cold 1X PBS, prior to 500 x g centrifugation at 4°C for 10 minutes and addition of 100 mL of ice-cold 10% DMSO (Sigma-Aldrich, cat #D2650), 10% FBS in RPMI 1640 to cell pellets. Cells were pipetted up and down twice and immediately stored at -80°C prior to delivery to the Epigenomics Development Laboratory at Mayo Clinic Rochester, where further preparation was completed. OMNI ATAC-seq was performed following a published protocol^58^. Amplified libraries were purified, and the sizes of library DNA were determined with a fragment analyzer (Advanced Analytical Technologies) using the High Sensitivity NGS Fragment Analysis Kit (Advanced Analytical Technologies, cat #DNF-486). The enrichment of accessible regions was determined by the fold difference between positive and negative genomic loci using real-time PCR. The libraries were sequenced to 51 base pairs from both ends on an Illumina NovaSeq 6000 SP instrument.

### Differential gene expression analysis for bulk RNA-sequencing

Analysis of sequencing data was carried out by bioinformaticians in the Quantitative Health Science department at Mayo Clinic Rochester. For bulk RNA-seq analysis, the raw RNA sequencing paired-end reads for the samples were processed through the Mayo RNA-Seq bioinformatics pipeline, MAP-RSeq (for human: v3.1.4, for mouse: v3.1.5)^59^. Briefly, MAP-RSeq employs the very fast, accurate and splice-aware aligner, STAR^60^, to align reads to the reference human genome build hg38 or the mouse genome build mm39. The aligned reads are processed through a variety of modules in a parallel fashion. Variants were called using the RVBoost module from MAPR-seq. Gene and exon expression quantification were performed using the Subread^61^ package to obtain both raw and normalized (FPKM – Fragments Per Kilobase per Million mapped reads) reads. Finally, FASTQC^62^, MultiQC^63^ and RSEQC^64^ tools were used for comprehensive quality control assessment of the aligned reads. Using the raw gene counts report from MAP-RSeq, differentially expressed genes between the groups were assessed using the bioinformatics package edgeR 2.6.2^65^ and reported using a level of significance set on fold change (log2 scale on x-axis of volcano plots, marked at log2(FC≥0.5) and p-value≤5% (log10 scale on y-axis of volcano plots, marked at -log10(p-value≤0.05)≥1.3).

### Differential gene expression analysis for bulk ATAC-sequencing

Analysis of sequencing data was carried out by bioinformaticians in the Quantitative Health Science department at Mayo Clinic Rochester. Bulk ATAC-seq data were processed to identify open chromatin regions (OCRs). Paired-end reads were aligned to the mouse genome (mm39) using Burrows–Wheeler Aligner (BWA, v0.5.9)^66^, and PCR duplicates were removed with Picard MarkDuplicates (v1.67). Peaks were called with MACS2 (v2.0.10)^67^ using default parameters. Differential accessibility analysis across experimental conditions was performed using the DiffBind package (v2.14.0)^68^, with significant OCRs defined as those with false discovery rate (FDR)≤0.05 and absolute log2(fold change>2). To assess transcription factor activity, footprinting analysis was performed using TOBIAS (v0.12.0)^69^, which corrects for Tn5 transposase sequence bias and infers occupancy at motif instances. Motif enrichment analysis of differentially accessible OCRs was conducted using the HOMER suite (v4.11), with enriched motifs ranked by significance. Peaks were visualized in the Integrative Genomics Viewer and aligned to Matched Annotation from NCBI and EMBL-EBI (MANE) transcripts.

### Single cell RNA sequencing and analysis

Mouse CD8^+^ TILs were isolated from adult mice (>8 weeks old) for single-cell RNA sequencing. Single cell libraries were prepared using Fluent Biosciences Pre-Templated Instant Partitions Sequencing (PIPseq) v3 Kit according to the manufacturer’s protocol. Briefly, cells were first filtered through a 40-μm cell strainer for the debris removal. The viability was assessed through Trypan Blue cell counting, and samples with more than 80% viability were proceeded for sample preparation. Each sample was loaded to the PIPseq microfluidic system, which partitioned individual cells into droplets containing molecular barcodes for unique transcript identification. During reverse transcription, unique molecular identifiers (UMIs) and well-specific barcodes were incorporated to enable precise quantification and tracking of individual transcripts. Subsequently, cDNA synthesis and amplification were performed to maximize transcript coverage and minimize bias. Amplified cDNA was purified and quantified using an Agilent 2100 Bioanalyzer. Following quality assessment, sequencing libraries were constructed by fragmenting cDNA, performing end-pair and A-tailing and ligating adapters compatible with the sequencing platform.

Sequencing was performed on an Illumina NovaSeq platform using the S4 flow cell platform in a paired-end format. Read 1 (28 bp) was used to sequence the transcript barcode, while Read 2 (91 bp) captured the transcript sequence. Unique Index read pairs (i7: 8 bp, i5: 8 bp) were used for demultiplexing. The sequencing was covered for an average depth of 40,000 reads per cell. Raw sequencing data were processed using Fluent Biosciences’ PIPseq analysis pipeline (v3.2). Reads were aligned to the GRCm39 reference genome, and unique molecular identifiers (UMIs) were used to deduplicate transcripts. Cells with fewer than 500 detected genes, greater than 20% mitochondrial transcript counts, or fewer than 1000 UMI counts were excluded from further analysis. Quality control and visualization were performed using Seurat (v4.3.0). Data were SCTransformed and scaled using Seurat. Highly variable genes were identified, and dimensionality reduction was performed using principal component analysis (PCA). The first 20 principal components were used to compute a uniform manifold approximation and projection (UMAP) for visualizing the dataset. Clustering was performed using the Louvain algorithm implemented in Seurat to identify groups of cells with similar expression profiles based on shared nearest neighbors in the principal component space. It was further optimized iteratively to capture biologically meaningful subpopulations across different scales. After initial clustering, sub-clustering was performed on individual groups to further resolve heterogeneous populations. The clusters and subclusters were visualized using UMAP to facilitate interpretation and identification of distinct cell populations. Differential expression analysis was performed to compare gene expression levels between clusters or predefined groups of cells. The FindMarkers function in Seurat was used with the Wilcoxon rank-sum test, a non-parametric method that evaluates differences in gene expression without assuming a normal distribution. For statistical significance, genes were considered differentially expressed if they had an adjusted p-value < 0.05 (calculated using the Benjamini-Hochberg method) and a log_2_fold change > 1. Data from multiple datasets were merged into a single data frame in R using the merge function, combining shared gene expression features. After integration, the merged dataset was normalized and scaled using Seurat to account for differences in sequencing depth and technical variability. Batch effects were corrected using Harmony (v1.0), which ensured effective alignment of datasets while preserving biologically meaningful variation.

### Statistical analysis

Statistical analysis for all assays, except for differential gene expression analysis from sequencing data and pathway analysis, was performed in GraphPad Prism (v10.1.1). Throughout the text statistical significance is designated as P*≤.05, P**≤.01, P***≤.001, and P****≤.0001. The definition for center and error for all bar graphs is the mean + SD or SEM. Individual points represent data from a single biological sample. The statistical test performed and the group size of n can be found in the corresponding figure legend.

## AUTHOR CONTRIBUTIONS

Conceptualization, AL, YL, HD; Data curation, AL, ED, JH, YL; Formal analysis, AL, ED, YL, HD; Funding acquisition, HD; Investigation, AL, JH, YL, JG; Methodology, AL, JH, JD, YL; Project administration, HD; Resources, HD; Software, YL; Supervision, HD; Visualization, AL, ED, JH, HD; Writing – original draft, AL, HD; Writing – review & editing, AL, JH, ED, YL, JG, HD.

## ACKNOWLEDGMENTS

We acknowledge all members of the Dong laboratory that contribute to scientific discussions, especially Xin (Cindy) Liu for technical help, and Susan M. Harrington for animal breeding. We acknowledge Dr. Alex Q. Wixom and Teja Koganti for bioinformatics analysis, carried out through the Quantitative Health Sciences department at Mayo Clinic Rochester. We also acknowledge the Genome Analysis Core for their support with sequencing, as well as the Epigenomics Program at Mayo Clinic Rochester for their help with ATAC-sequencing preparation. We utilized BioRender.com to generate graphics found throughout the manuscript.

